# Self-organized spatial targeting of contractile actomyosin rings for synthetic cell division

**DOI:** 10.1101/2024.06.17.599291

**Authors:** María Reverte-López, Nishu Kanwa, Yusuf Qutbuddin, Marion Jasnin, Petra Schwille

## Abstract

One of the challenges of bottom-up synthetic biology is the engineering of a minimal module for self-division of synthetic cells. To produce the contractile forces required for the controlled excision of cell-like compartments such as giant unilamellar vesicles (GUVs), reconstituted cytokinetic rings made of actin are considered to be among the most promising structures of a potential synthetic division machinery. Although the targeting of actin rings to GUV membranes and their myosin-induced constriction have been previously demonstrated, large-scale vesicle deformation has been precluded due to the lacking spatial control of these contractile structures. Here, we show the combined *in vitro* reconstitution of actomyosin rings and the bacterial MinDE protein system, effective in targeting *E.coli* Z-rings to mid-cell, within GUVs. Incorporating this spatial positioning tool, which induces active transport of any diffusible molecule on membranes, yields self-organized assembly of actomyosin rings at the equatorial plane of vesicles. Remarkably, the synergistic effect of Min oscillations and the contractile nature of actomyosin bundles induces mid-vesicle membrane deformation and striking bleb-like protrusions, leading to shape remodeling and symmetry breaking. Our system showcases how functional machineries from various organisms may be synergistically combined *in vitro*, leading to the emergence of new functionality towards a synthetic division system.

## 1. Introduction

Bottom-up synthetic biology is an interdisciplinary field currently fostering promising technological advancements to tackle the environmental and biomedical challenges of the future while it strives towards a fundamental goal: the construction of an artificial cell from a set of minimal functional modules.^1–4^ As protocell models, the biomimetic chassis commonly used in the field are giant unilamellar vesicles (GUVs), membrane-enclosed containers capable of hosting biochemical reactions.^5^ To ensure the autonomy and continuity of our artificial vesicular systems, key cellular features and processes must be recapitulated within protocells; particularly, their ability to divide and self-replicate, a critical step in a cell’s life cycle.^1,6^

In this regard, several strategies have been conceived to engineer a synthetic division module capable of mechanical membrane abscission.^7^ While the reconstitution of well-characterized bacterial divisome machinery is a very promising approach,^8,9^ recapitulating division with a eukaryotic cytoskeleton-based toolbox is another intriguing strategy, due to the versatility and modularity of eukaryotic division proteins. Inspired by eukaryotic cytokinesis, the so-called engineering route works on assembling an actomyosin contractile ring at the GUV equator which, upon its controlled diameter reduction, is supposed to constrict the vesicle membrane until scission.^10^ Two main cytoskeletal components are presumably required for this ring assembly route: actin and myosin. Actin, in its filamentous form and together with its many regulatory binding proteins, assembles into bundles which positioned at mid-cell constitute the ring scaffold. To generate the tension required for ring constriction, actin must associate with myosin, the key motor protein that drives contractility of the actin assembly and induces furrow ingression.^11,12^

Several studies have shown the successful reconstitution of contractile actomyosin rings *in vitro*.^13–15^ Of particular interest is the reconstitution inside GUVs by Litschel *et al*.^14^ Using talin-vinculin as bundlers, membrane-bound actomyosin rings induced transient deformation in vesicles. However, bundles slipped on the membrane and formed condensate clusters, impeding the radial targeting of contractile forces on the vesicle membrane. Indeed, a key requirement for the cytokinetic engineering route to succeed is the stable circumferential positioning of the contractile ring during constriction of the vesicle, ideally at mid-cell. To date, however, spatiotemporal control of actomyosin rings in GUVs has not been achieved. For the eukaryotic reconstitution approach this challenge is of particular complexity, since reconstitution of *in vivo* mechanisms to maintain mid-cell ring placement would require the synergistic integration of many proteins systems and signalling molecules, a currently unattainable feat.^10^

Interestingly, recent studies have shown the robustness and versatility of a protein-based spatial positioning toolset: the MinDE system.^16–18^ The *Escherichia coli* (*E. coli*) Min proteins are a reaction-diffusion system able to self-organize on membranes through ATP hydrolysis. Composed of three proteins – MinD, MinE and MinC – the Min complex has a particular function *in vivo*: the localization of the FtsZ division ring (Z-ring) in the middle of rod-shape bacteria. Their self-organization mechanism consists of three steps: (1) ATP-dependent dimerization of MinD promoting membrane attachment, (2) MinE recruitment to membrane-bound MinD stimulating MinD’s ATP-ase activity, (3) ATP hydrolysis and detachment of the MinDE complex from the membrane. Following this mechanism of pattern formation, MinD and MinE oscillate from one pole of the cell to the other and inhibit the assembly of the Z-ring near the poles via depolymerization of FtsZ through MinC, the functional cargo protein that is not involved in MinDE self-assembly.^19^ *In vitro*, however, a surprising functionality of the MinDE system was discovered.^20^ When reconstituted on planar supported lipid bilayers and inside GUVs, the proteins MinD and MinE can non-specifically sort and position any membrane-bound cargo. More precisely, through a diffusiophoretic transport mechanism,

MinD fluxes can spatiotemporally control molecules on the membrane by frictional forces and generate anti-correlated molecular patterns.^16^ This newly discovered biochemical function, although possibly irrelevant *in vivo*, could thus be exploited for the positioning of membrane-attached molecules and other biomimetic features in artificial systems.^17,18^

In this study, we demonstrate the successful co-reconstitution of actomyosin architectures and the MinDE system inside GUVs. Upon optimization of encapsulation conditions, time-lapse imaging revealed the MinDE-driven diffusiophoretic positioning of actomyosin rings and bundle networks at mid-cell, where we observed equatorial furrow-like invaginations breaking spherical symmetry. Moreover, we show that, besides the spatiotemporal control of actomyosin bundles at the membrane, MinDE binding can induce bleb-like outward protrusions in single-phase vesicles and domain-specific deformations in phase-separated GUVs. Thus, the experimental insights here reported demonstrate that upon ATP hydrolysis, MinDE proteins not only aid in active contractile ring localization, but also generate mechanical work to remodel vesicle membranes during synthetic division. Overall, these results showcase the advantages of integrating synthetic toolsets of different origin to engineer artificial cells with advanced functionalities.

## 2. Results

### 2.1. Co-reconstitution of actomyosin networks and the MinDE system inside GUVs

To achieve the co-reconstitution of actomyosin networks and Min oscillations inside vesicles, we carried out encapsulation experiments via double emulsion transfer to identify optimal experimental conditions for the dynamic and functional interplay of both systems’ components.

First, since G-actin and Min protein self-organization depend on factors like salt concentration, supply of ATP and the presence of divalent cations in solution, we tuned the inner environment of the vesicles to simultaneously facilitate actin filament polymerization and MinD dimerization, critical for MinD interaction with negatively charged amphiphiles and its cooperative binding to membranes. Given the importance of the membrane as catalytic matrix for the spatiotemporal organization of MinDE proteins, we generated vesicles containing negative charge in the bilayer to enable the self-organization of Min proteins into different oscillation modes.^21,22^ In addition, we incorporated biotinylated lipids to link biotinylated actin filaments to the inner leaflet of the GUVs. Anchoring actin assemblies to the membrane through neutravidin-biotin bonds allowed us to exploit the diffusiophoretic capabilities of Min proteins, which require membrane-bound cargo to induce the ATP-driven transport of molecules on membranes.^16^

Once optimal buffer conditions and membrane composition were identified, we chose fascin as the crosslinking protein to generate high-order actin bundle structures. As previously reported,^14^ by binding fascin-assembled bundles to the membrane via neutravidin-biotin bonds we obtained long and curved bundles that robustly bound to the membrane adapting to the vesicle curvature. Additionally, to accelerate actin polymerization kinetics and decrease MinDE wavelength and oscillation velocity, we employed Ficoll70 as macromolecular crowder,^23,24,9^ which also facilitated vesicle production. Finally, to make our actin assemblies contractile and render membrane deformations,^13,14^ the motor protein myosin II was added to the inner solution mix (**Fig. 1a**).

**Fig. 1.**
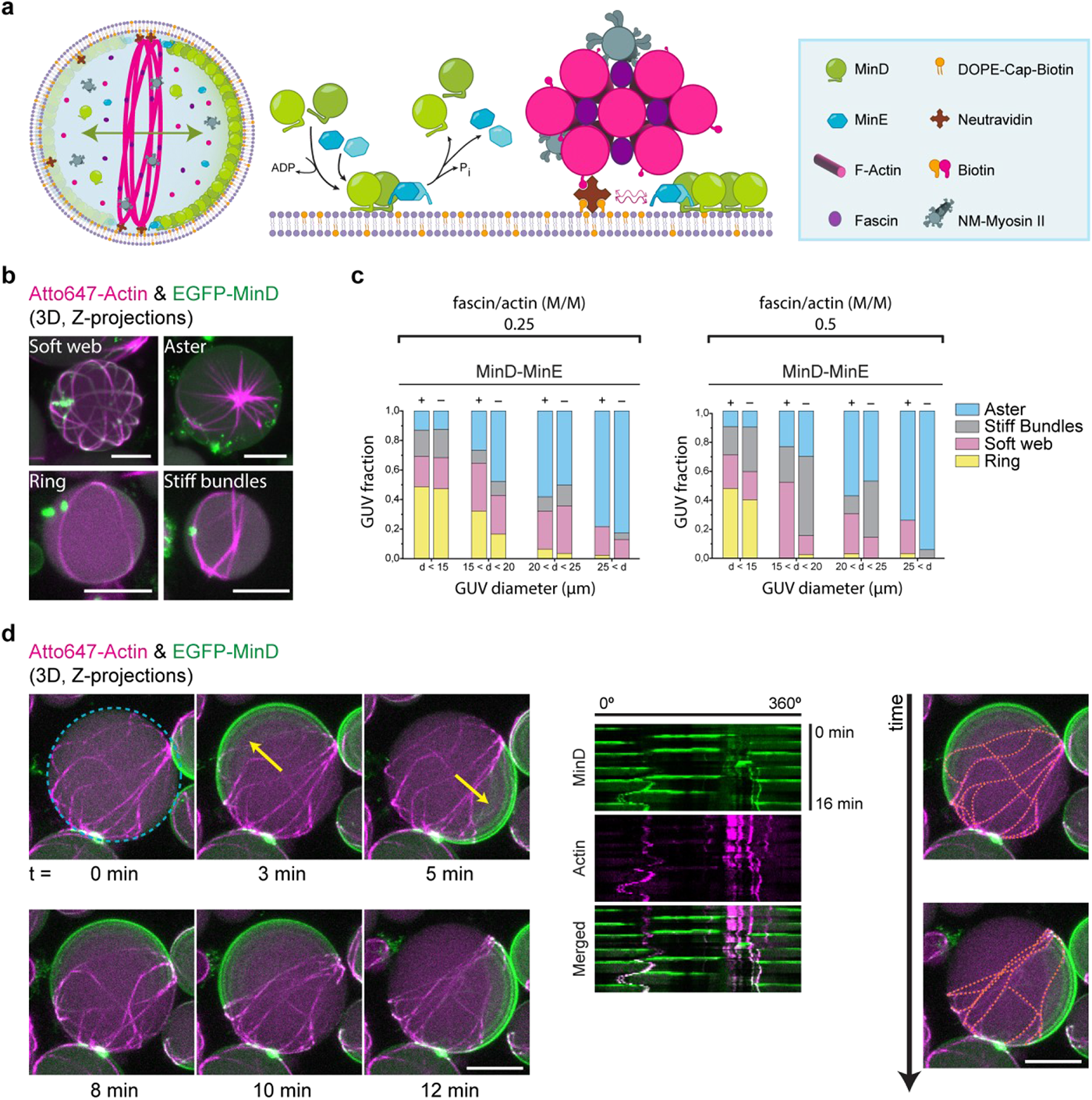
Co-reconstitution of actomyosin networks and the MinDE system enables the reorganization and positioning of actomyosin bundles at mid-cell. **a** Schematic illustration of the GUV content and the two macromolecular reactions at membrane level: the MinDE self-assembly mechanism behind pattern formation and the diffusiophoresis-mediated transport of neutravidin-bound actomyosin bundles by Min proteins. The active flux of MinDE proteins on the vesicle membrane interacts non-specifically via frictional forces with membrane-bound neutravidin inducing the transport and positioning of these molecules, and consequently the actomyosin bundles linked to them, towards areas of low MinD density. **b** 3D projections of confocal images showing the 4 phenotypes of actin architectures obtained after encapsulating 2.4 µM actin, 0.6 µM fascin (fascin/actin molar ratio = 0.25), 0.05 µM myosin II, 50 g/L Ficoll70, 3 µM MinD, 3 µM MinE and 5 mM ATP. Scale bars: 10 µm. **c** Bar graphs with the frequencies of the four actomyosin phenotypes observed at different vesicle diameters when encapsulation experiments were performed at 0.25 and 0.5 fascin/actin molar (M/M) ratio in the presence and absence of Min proteins and protein/crowding conditions specified in b. Experiments performed per condition n = 3, total number of GUVs analysed per condition = 150. **d** 3D projections of time-lapse confocal images depicting the reorganization and stacking of actomyosin bundles towards the vesicle equator driven by the diffusiophoretic transport of Min pole-to-pole oscillations. Yellow arrows indicate the perpendicular orientation of MinDE oscillations with respect to actomyosin bundles, which get antagonistically positioned at mid-cell. Kymographs generated at the vesicle equator (blue dashed circle) are meant to visually define the position of fluorescent features at this region over time. Orange dotted lines depict the approximate distribution of actin bundles on the membrane at two time points. Vesicle content as specified in b. Scale bars: 10 µm.

To investigate the effects of MinDE oscillations on the formation of actin-bundle architectures under crowding conditions, we encapsulated actin and fascin at different molar ratios together with myosin II in the presence and absence of Min proteins. To this end, we analysed actin architecture types and quantified their frequency of appearance in terms of GUV sizes. Similar to recent cytoskeletal reconstitutions in GUVs,^15,25^ we observed four main actin phenotypes in both the presence and absence of Min proteins: soft bundle webs, actomyosin asters, flexible rings and stiff-straight bundles (**Fig. 1b**). Although phenotype yields differed, Min oscillations supported the bundling and assembly of actomyosin architectures on GUV membranes. In particular, at a 0.25 fascin/actin molar ratio (**Fig. 1c**), when Min proteins were part of the reaction mix, we detected an increase in flexible ring yields, as well as a lower probability of stiff bundle formation irrespective of vesicle size. As previously reported, confinement impacts actin-bundle architecture.^13,25^ In both samples containing or in absence of Min proteins, the probability of flexible ring formation was significantly higher in small diameter vesicles (diameter < 15 µm), whereas for medium and big vesicles the predominant phenotype was aster, reaching almost 50% formation probability in vesicles between 20-25 µm in diameter and 80% for vesicles bigger than 25 µm. Similarly, when a 0.5 fascin/actin molar ratio was employed and a higher fascin/actin concentration was encapsulated, we observed the four aforementioned phenotypes in the presence and absence of Min proteins (**Fig. 1c**). Under these conditions, however, only the frequency of stiff bundle formation decreased upon addition of Min proteins and the probability of flexible ring formation drastically decreased for GUVs bigger than 15 µm diameter in samples containing Min proteins or in their absence.

Taken together, we show the successful co-reconstitution of the actomyosin system together with Min proteins and demonstrate that MinDE oscillations are compatible with the assembly of membrane-bound actomyosin architectures inside GUVs, and vice versa. Notably, addition of Min proteins promotes the formation of flexible actomyosin rings in all vesicle sizes encapsulated with a 0.25 fascin/actin ratio.

### 2.2. Diffusiophoresis-mediated positioning of actomyosin bundles at mid-cell by the MinDE system

Having established the conditions to reconstitute dynamic Min oscillations together with actomyosin-bundle assemblies inside GUVs, we then investigated whether Min proteins could effectively reorganize these assemblies and position them at mid-cell via their diffusiophoretic mechanism of molecular transport. Since flexible rings and soft bundle webs are the two types of actin architectures that could efficiently transmit contractile forces to the membrane, we studied the spatiotemporal organization of these two phenotypes by Min oscillations with time-lapse microscopy.

In agreement with past studies,^26^ we observed three main Min oscillation modes resultant from the reaction-diffusion fluxes of Min proteins on the inner leaflet of vesicles (**Supplementary Fig. 1a**): pulsing (oscillation characterized by the consecutive binding and unbinding of MinD to the entire vesicle membrane), pole-to-pole (sequential binding of MinD to the hemispheres of the vesicle), and circling waves (MinDE waves revolving around the inner leaflet of the membrane).

As MinDE pole-to-pole oscillations are the desired phenotype to actively transport molecules to the mid-cell region via diffusiophoresis,^9,20^ we first scrutinized actomyosin-containing vesicles exhibiting this dynamic pattern. Strikingly, in vesicles containing actomyosin bundles isotropically distributed all over the membrane, MinDE pole-to-pole oscillations yielded an anticorrelated and directional movement of the bundles perpendicular to the oscillation axis, reducing bundle interdistance and accumulating them at mid-cell (**Fig. 1d, Supplementary Movie 1**). Subjected to the highly dynamic MinDE pattern, the actomyosin bundles still showed positional fluctuations at the GUV equator over time, but maintained a perpendicular orientation to the oscillation axis.

Subsequently, to test the robustness of the MinDE diffusiophoretic transport in our actin-based encapsulation system, we varied the experimental conditions from our standard inner solution mix. We found that, in the absence of myosin II, under varying Ficoll70 concentrations (10-50 g/L), and employing different molar ratios for fascin/actin (0.25 or 0.5) as well as MinD/MinE ratios (1 or 2), Min proteins were capable of actively arranging membrane-bound actin bundles via diffusiophoresis when patterns different from pulsing developed on the vesicle membrane. Importantly, and as expected from our previous experiments, MinDE pole-to-pole patterns rotated and positioned fascin-assembled actin rings in the absence of myosin II, maintaining ring orientation perpendicular to MinDE oscillations (**Supplementary Fig. 2b**). In addition, when MinDE circling patterns emerged on vesicles containing fascin-bundled actin rings, we observed that the frictional forces induced by the directional MinD protein flux promoted the circular displacement of one end of the ring towards the opposite end (**Supplementary Fig. 2c**), resulting in the complete folding of the ring in less than 15 minutes (**Supplementary Fig. 2d**). Moreover, we detected that chaotic MinDE patterns – a dynamic mode in which Min proteins bind and unbind membrane areas in stochastic direction – could also alter the distribution of actomyosin bundles and, in some instances, buckle and collapse the network (**Supplementary Fig. 1b,c**).

Thus, our data demonstrate that the MinDE system can be used to regulate the spatiotemporal localization of membrane-bound actomyosin bundles and most importantly, position contractile actomyosin architectures at mid-cell via its characteristic pole-to-pole oscillation mode.

### 2.3. Equatorial constriction of vesicles induced by positioned actomyosin architectures

In our system, myosin II not only acts as a crosslinking agent but also provides the contractile force required to induce membrane deformations. Having shown that the MinDE system localized actomyosin assemblies at the vesicle equator, we next investigated whether the positioned contractile assemblies could generate furrow-like membrane deformations.

First, to explore the contractile effect of myosin II on positioned actin structures, we carried out encapsulation experiments at 0.25 and 0.5 fascin/actin molar ratio with 50 g/L Ficoll70 and examined the vesicles presenting pole-to-pole oscillations with time-lapse confocal microscopy. Interestingly, when we employed a 0.5 fascin/actin molar ratio, mid-cell deformation induced by actomyosin ring constriction could be observed (**Fig. 2a**). This furrow-like invagination of the membrane, sustained over time (**Supplementary Movie 2**), generated two lobes where MinD proteins continued oscillating in a pole-to-pole pattern maintaining the localization of the ring at mid-cell.

**Fig. 2.**
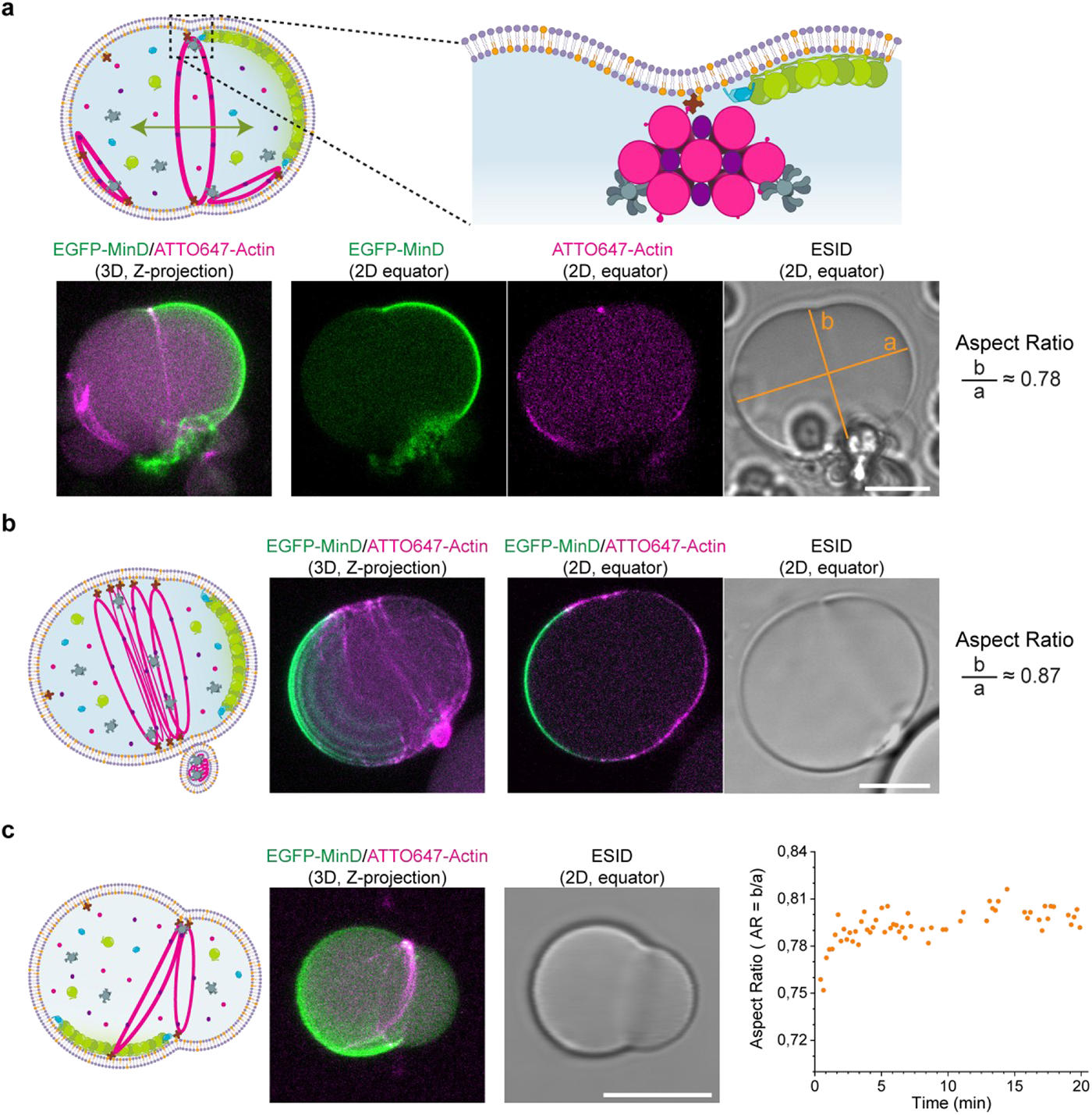
Positioned actomyosin rings and soft webs constrict vesicles at mid-cell. **a** Schematic illustration behind the mechanism of membrane deformation. Contractile actomyosin bundles positioned by MinDE proteins at mid-cell induce furrow-like membrane invaginations. 3D projections and 2D confocal images show an actomyosin ring constricting the vesicle at its equator. Orange lines indicate the major (a) and minor (b) axes measured to calculate the aspect ratio of the deformed vesicle (for spherical vesicles: aspect ratio = 1). Inner solution mix: 4 µM actin, 2 µM fascin (fascin/actin molar ratio = 0.5), 0.05 µM myosin II, 50 g/L Ficoll70, 3 µM MinD, 3 µM MinE and 5 mM ATP. Scale bar: 10µm. **b** Schematic illustration, 3D projections and 2D confocal images of a vesicle containing a soft web of actomyosin bundles at the vesicle centre being positioned by pole-to-pole Min oscillations. The contractile actomyosin band formed causes the deformation of the vesicle (aspect ratio < 1). Inner solution mix: 2.4 µM actin, 0.6 µM fascin (fascin/actin molar ratio = 0.25), 0.05 µM myosin II, 50 g/L Ficoll70, 3 µM MinD, 3 µM MinE and 5 mM ATP. Scale bar: 10µm. **c** Schematic illustration, 3D projection and 2D confocal image of a vesicle with a non-positioned contractile actomyosin assembly due to the loss in pole-to-pole MinDE oscillations. Constriction of the actomyosin bundles results in the deformation of the vesicle membrane into an asymmetric dumbbell shape. Scatter plot depicts the aspect ratio of the vesicle at different time points. Inner reaction mix: 4 µM actin, 2 µM fascin (fascin/actin molar ratio = 0.5), 0.05 µM myosin II, 20 g/L Ficoll70, 3 µM MinD, 3 µM MinE and 5 mM ATP. Scale bar: 10 µm.

In line with these observations, we also detected vesicle deformation at 0.25 fascin/actin molar ratio. Contrary to single rings, soft actomyosin bundle networks positioned at the vesicle equator formed a constriction band resulting in the loss of spherical vesicle shape (**Fig. 2b**). In addition to this large-scale membrane deformation, we observed that an actin-filled membrane out-bud formed at the constriction site (**Supplementary Movie 3**). Litschel *et al.* already reported on this type of membrane deformation, which results from the sliding of actomyosin bundles along the membrane to one side of the vesicle and their collapse into a condensate.^14^ Notably, the membrane bud we observed was localized closely to the actomyosin band constricting the vesicle at the equator, while MinDE pole-to-pole oscillations continued at the two hemispheres generated at each side of the actomyosin band. To quantitatively assess vesicle deformation, we calculated the vesicle aspect ratio (AR) as the ratio between their major (a) and minor (b) axes. In both examples presented, aspect ratios indicated a vesicle deformation towards a rod-like shape (AR < 0.9).

Furthermore, given that macromolecular crowders can impact both actin bundle architecture and the mechanical properties of the vesicle membrane,^27,28^ we subsequently performed encapsulation experiments at lower crowding concentration (20 g/l Ficoll70) and found membrane deformations in line with those already showed. Furthermore, under these experimental conditions, we detected an example of eccentric membrane deformation caused by an actomyosin ring when the established MinDE oscillation pattern inside the vesicle was different from the pole-to-pole mode (**Fig. 2c**). More specifically, the MinDE pattern at the vesicle membrane transitioned into a circling and less dynamic MinDE oscillation (possibly due to ATP depletion). Contrary to vesicles presenting mid-cell constriction, the membrane deformation observed induced a characteristic asymmetric dumbbell shape with two differently sized sub-compartments. Nevertheless, this asymmetric deformation, sustained over time (average AR over 20 minutes = 0.79), did not collapse after more than an hour of imaging (**Supplementary Movie 4).** Myosin II and ATP concentration added were 0.05 µM and 5 mM, respectively. Further experiments are therefore required to find the optimal encapsulating conditions enabling MinDE-stabilized rings positioned at mid-cell to undergo progressive contraction and controllably reduce their diameter by the action of myosin II motors.

In summary, we show that MinDE pole-to-pole oscillations can target the constriction of actomyosin architectures at the vesicle equator, resulting in the generation of sustained furrow-like membrane deformations.

### 2.4. MinDE-induced blebbing in reconstituted actomyosin vesicles

When thicker and more abundant actomyosin bundles developed at the membrane in the form of soft webs, we detected the establishment of more chaotic and static MinDE patterns. Thus, to further characterize the system, we performed time-lapse imaging on vesicles presenting chaotic or static-like MinDE patterns.

Interestingly, a large number of vesicles exhibiting these patterns developed membrane deformations similar to bleb-like morphologies (**Fig. 3a, Supplementary Movie 5**). To get further insights into the underlying mechanism behind bleb formation, we analysed the interaction between our co-reconstituted protein systems and the vesicle membrane. Closer inspection of the blebs’ cross-sections revealed that the actomyosin soft web inside the vesicle compartmentalized the membrane into areas delimited by the peripheral attachment of bundles (**Fig. 3b, Supplementary Fig. 3a**). Due to this compartmentalization, container symmetry was lost and MinDE proteins generated a chaotic oscillation capable of deforming the membrane. More specifically, we found that the initial spontaneous curvature induced on these compartments by MinDE binding increased as more MinD molecules were recruited to the membrane, resulting in the outward growth of the dynamic bleb-like protrusions (**Fig. 3c, Supplementary Fig. 3b**). Subsequently, after MinDE-driven membrane deformations, we observed that the reduction in bilayer tension and the recovery of the initial vesicle shape was accompanied by the generation of a membrane out-bud (**Fig. 3a**, blue arrows), which was not reabsorbed into the mother vesicle (**Supplementary Fig. 3c**).^29^

**Fig. 3.**
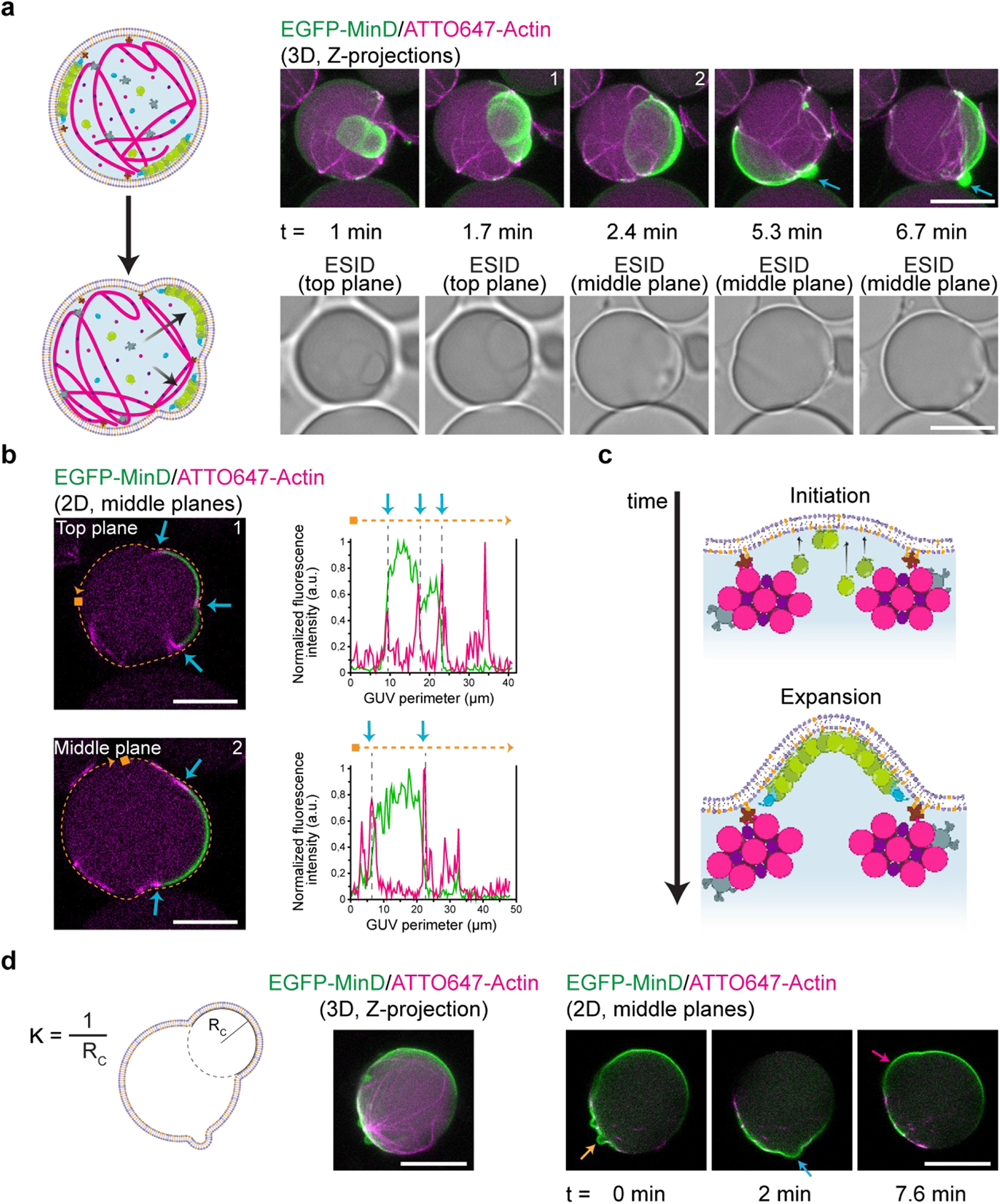
MinDE-induced blebbing in vesicles containing reconstituted actomyosin architectures. **a** Schematic illustration depicting the change in vesicle shape due to MinDE chaotic oscillations. Min proteins attach to areas delimited by soft actomyosin bundles and deform the membrane generating dynamic bleb-like protrusions. Fluorescence and brightfield confocal time-series show a blebbing vesicle. After bleb retraction, the reduction in bilayer tension generates an outward lipid bud (blue arrows). Encapsulation conditions: 2.4 µM actin, 0.6 µM fascin, 0.05 µM myosin II, 50 g/L Ficoll70, 3 µM MinD, 3 µM MinE and 5 mM ATP. Scale bars: 10 µm. **b** Confocal cross-section images at two time points of the vesicle in section a. Peripheral actomyosin anchoring creates a delimiting area which deforms upon MinDE binding. Additionally, MinDE diffusiophoretic transport changes the position of actomyosin bundles and the shape of the membrane area available for Min protein recruitment (blue arrows). Fluorescence intensity line plots of EGFP-MinD (green) and ATTO647-actin (magenta) demonstrate the demixing of both protein systems at the membrane perimeter (orange dotted line). Scale bars: 10 µm. **c** Schematic illustration of the proposed mechanism behind MinDE-induced blebbing. The recruitment of MinDE proteins to the compartmentalized inner leaflet of the bilayer generates the effect of a membrane outward protrusion in bleb form. **d** Schematic illustration depicting the radius of curvature R_C_ used to calculate the curvature (Κ = 1/R_C_) of the blebs. 3D projection and 2D time-lapse confocal images show a vesicle with diverse bleb-like deformations emerging over time. Orange arrow points at a bleb with Κ = 0.73 µm^-1^. Blue arrow, Κ = 0.27 µm^-1^. Magenta arrow, Κ = 0.10 µm^-1^. Encapsulation mix: 4 µM actin, 2 µM fascin, 0.05 µM myosin II, 50 g/L Ficoll70, 3 µM MinD, 3 µM MinE and 5 mM ATP. Scale bars: 20 µm.

Furthermore, consistent with our previous observations and simultaneous to this membrane remodelling effect, the diffusiophoretic transport of actomyosin bundles reorganized the network at the membrane. As a result, membrane compartments changed in size and the oscillations maintained a chaotic mode inducing dynamic blebs in other areas of the vesicle (**Fig. 3b**).

To further scrutinize the membrane-remodelling capabilities of Min proteins along with our actomyosin architectures, we performed encapsulation experiments at 0.5 fascin/actin molar ratio and calculated the curvature of the blebs observed. Similar to our non-deflated vesicles encapsulated with 0.25 fascin/actin ratio, MinDE binding induce blebbing in a subset of vesicles (**Supplementary Movie 6**). Time-lapse imaging revealed that Min proteins can induce blebs with a wide range of curvatures (Κ, calculated as the inverse of the radius of a circle that fits the bleb). As MinDE established a chaotic oscillation inside the vesicle, small blebs (Κ = 0.73 µm^-1^), medium size (Κ = 0.27 µm^-1^) and big (Κ = 0.10 µm^-1^) deformations emerged (**Fig. 3d**, orange, blue, and magenta arrows, respectively).

Hence, our results show that, when co-reconstituted with actomyosin bundle networks, the MinDE system can generate dynamic bleb-like outward protrusions in vesicles encapsulated at iso-osmolar conditions, confirming its capabilities as a membrane remodeling protein system as previously observed.^26,30^

### 2.5 Co-reconstitution of actomyosin bundle networks and the MinDE system in phase-separated vesicles show remodelling of membrane domains

Lastly, to increase the complexity of the system and test its compatibility with other shape remodeling strategies, we tested its reconstitution in vesicles of ternary lipid mixtures. Due to their tuneable mechanical and biochemical properties, phase-separated lipid membranes constitute another approach to aid in the remodeling of biomimetic systems by altering membrane curvature, fluidity, etc.^31–33^ Furthermore, two-phase vesicles constitute an additional strategy to study the reorganization and deformation of free-standing lipid domains by actomyosin networks.^34–36^

To achieve the co-reconstitution of the two protein systems in GUVs with phase-separated lipid domains we again employed the double emulsion transfer method. At room temperature (25 °C), GUVs demixed into coexisting Liquid-ordered (Lo) and Liquid-disordered (Ld) domains, where Ld domains consisted of DOPE-Biotin to facilitate actin binding (**Fig. 4a**). To study the successful reconstitution of the system inside phase-separated vesicles, we again performed time-lapse confocal imaging and observed that Min proteins could oscillate by binding to Ld domains on the vesicle membrane. Notably, due to the high frequency of soft actomyosin bundle webs formed, MinDE proteins also established chaotic oscillations on the Ld domains of the vesicle. The flexible bundles, however, spanned and crossed both lipid domains.

**Fig. 4.**
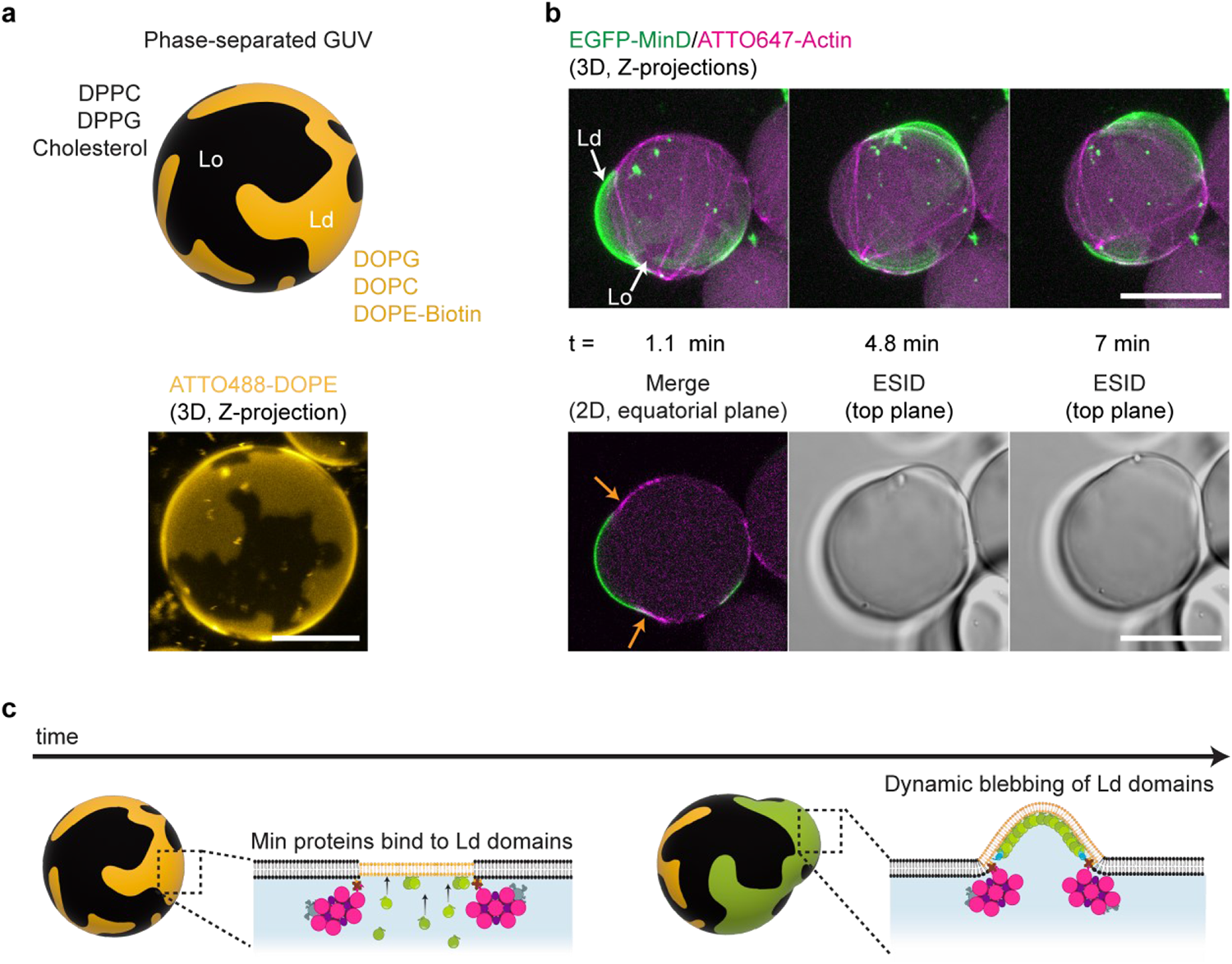
MinDE-induced bleb morphologies on phase-separated GUVs with encapsulated actomyosin architectures. **a** Schematic illustration (top) and 3D confocal image (bottom) show the membrane composition employed to generate phase-separated vesicles and the domains obtained. Scale bar: 10 µm. **b** 3D projections and 2D confocal images depict a blebbing phase-separated vesicle. MinDE proteins bind and oscillate on Ld domains. Actomyosin bundles remain at lipid-phase boundaries as Min proteins transiently deform Ld domains (orange arrows). Inner encapsulation mix: 2.4 µM actin, 0.6 µM fascin (fascin/actin molar ratio = 0.25), 0.05 µM myosin II, 20 g/L Ficoll70, 3 µM MinD, 3 µM MinE and 5 mM ATP. Scale bars: 20 µm. **c** Schematic illustration of the proposed mechanism behind the dynamic deformation of Ld domains by MinDE protein oscillations.

Interestingly, MinDE-driven diffusiophoretic transport displaced the actomyosin bundles bound at Ld/Lo boundaries, reorganizing the actomyosin network inside the vesicle. Moreover, consistent with our previous experiments performed on single-phase GUVs, MinDE binding to areas delimited by actomyosin bundles at domain boundaries deformed Ld domains into dynamic bleb-like protrusions (**Fig. 4b, Supplementary Movie 7**). In contrast to studies in which bulging or budding of phase-separated GUVs is externally induced by tuning membrane composition or changing osmotic conditions, and where deformations arise by an imbalance between surface tension and interfacial line tension,^33,37^ our phase-separated GUVs remained spherical over time in the absence of Min proteins, and no deformations in the form of blebbing or budding were observed (**Supplementary Fig. 4, Supplementary Movie 8**). Only in the presence of MinDE oscillations in GUVs containing actomyosin networks, timelapses showed the active – i.e., energy-consuming – deformation of Ld domains into outward blebs (**Fig. 4c**). During the course of these transient outward deformations, we also observed the remodeling of domains at the membrane such as their manoeuvring followed by splitting (**Supplementary Movie 7**). As a result of MinD binding to Ld domains, the initial demixing of lipid phases on the vesicle thus changed, and domain reorganization occurred on the entire vesicle. Interestingly, while the resultant dynamic protrusions in the membrane comprised of Ld domains, Lo domains stayed intact.

In conclusion, we demonstrate that the co-reconstitution of Min proteins and actomyosin can deform complex ternary vesicle membranes and dynamically remodel phase-separated lipid domains by rearranging and reshaping them on the membrane.

## 3. Discussion

In this study, we have successfully co-reconstituted contractile actomyosin rings and other assemblies with a protein-based spatial positioning toolset, the MinDE system. Our results demonstrate that, under optimal encapsulating conditions, actomyosin bundles can be spatiotemporally controlled at the membrane and positioned at mid-cell via pole-to-pole MinDE oscillations. Notably, the positioned bundles, due to their contractile nature, induced membrane deformations, breaking GUV spherical symmetry and generating furrow-like invaginations. In addition to the observed actomyosin-driven membrane remodeling, we provide direct evidence that MinDE proteins generate dramatic bleb-like deformations upon dynamic binding to the membrane, another source of symmetry breaking for the GUVs. Thus, our findings lend further credence to the hypothesis that the diffusiophoretic function of the MinDE system can be exploited to maintain actomyosin ring localization at mid-cell, one of the key milestones so far not successfully reported in the eukaryotic-based approach to synthetic cell division.

It should be noted that, after the adjustment in encapsulating conditions, our reconstitution conclusively showed the robustness of the MinDE system when integrated with the contractile actomyosin toolset. Not only did we observe the main MinDE-oscillation phenotypes extensively reported in past studies,^9,26^ our results also indicate that the establishment of Min patterns on the membrane had no detrimental effect of the formation of lipid-linked actomyosin structures. Indeed, we observed an increase in the frequency of GUVs containing membrane-bound rings when Mins were part of the inner encapsulation mix. Moreover, in agreement with previous MinDE geometry-sensing studies and the spatiotemporal feedback observed with FtsZ,^38,39^ our results demonstrate that Min oscillations orient themselves in a pole-to-pole fashion perpendicular to the spatially positioned ring structures while vesicles acquired ellipsoidal shape due to actomyosin contraction, an important aspect to consider as constriction progresses.

Alternative strategies for ring localization, like the use of microfluidic traps and curvature inducing/sensing biomolecules (e.g., septins, BAR domains, DNA origami), propose the use of external mechanical forces or biomolecules to generate a furrow-like negative curvature that could favour ring assembly.^40–44^ However, due to progressive membrane deformation, ring alignment could be lost.^40^ The MinDE system, by contrast, dynamically responds to vesicle shape changes, as MinDE pole-to-pole oscillations block any membrane binding on the poles, thereby enabling the targeting of mechanical forces at the centre.^9^ Nonetheless, to keep the actomyosin ring in place and prevent slippage of the bundles on the membrane, future research should investigate the co-reconstitution of scaffold (e.g., anillin, septins) and severing/depolymerizing proteins to increase turnover dynamics within actin bundles at the membrane as myosin contraction remodels the ring.^45,46^

Besides accomplishing the active positioning of actomyosin architectures, time-lapse imaging revealed a surprising new finding, the MinDE-driven blebbing of vesicles. *In vivo*, blebbing occurs at the cell poles as a mechanism to reduce cortical tension and ensure the stability of cellular shape during eukaryotic cytokinesis.^47^ In our *in vitro* reconstitution, however, blebs originate because of a mechanical MinDE function unfolded on free-standing membranes, and constitute a source of asymmetric remodeling of the membrane.^48^ While past studies already showed shape fluctuations in vesicles due to dynamic MinDE pulsing patterns at hypertonic and (near)-isotonic conditions,^26,30,49,50^ we hypothesize that the dramatic bleb-like protrusions here shown are due to confinement effects induced by actomyosin bundles creating lateral diffusion barriers for Min proteins, hence altering MinDE membrane kinetics and pole-to-pole patterns.^9^ Consequently, vesicle deformations could be attributed to the previously reported interactions and effects Min proteins have on membranes, such as: Min-induced increase of local viscosity,^51^ the generation of bilayer asymmetry due to the insertion of their alpha-helix at the inner leaflet,^48–50^ or even a possible scaffolding effect as a result of their oligomerization into flexible linear or 2D polymers.^52,53^ Although the change in spontaneous membrane curvature by MinDE oscillations requires further investigation, this compelling mechanical force driven by ATP hydrolysis could be exploited at the lateral regions of the vesicle for shape-symmetry breaking during synthetic division.

Furthermore, we observed an interesting mechanical effect on actin bundles themselves, arising via diffusiophoresis from MinDE dynamic patterns like chaotic or circling oscillations. Interestingly, time-lapse imaging showed bundles bending, collapsing, and stretching due to the diffusiophoretic frictional force of MinD fluxes. Thus, the system here reconstituted presents itself as another approach to study the crosslinker-dependent mechanical properties of actomyosin bundles and investigate the extent of diffusiophoresis when stiff supramolecular structures are bound to the membrane.

Importantly, and in accordance with previous studies, we observed actomyosin-driven deformation of GUVs under isosmotic conditions towards both ellipsoidal and asymmetric lobe shapes.^14,15^ However, after the stalling of myosin II on bundles occurred, contraction was arrested and deformation did not progress. It should be borne in mind that, in our system, myosin II also crosslinks actin and consumes ATP, an energy source employed by actin as well as MinD. Thus, we consider it reasonable to suggest that under our encapsulating conditions with fascin as bundling agent, myosin II contracts the actin assembly until it reaches full condensate formation or stalls due to a compacted actin architecture or ATP shortage.^27,54^ To effectively achieve a controlled contraction of the ring and its decrease in diameter on the membrane, there are a few approaches that could be implemented in our system. From the activation of latent myosin II with blebbistatin to trigger contraction late after encapsulation,^55,56^ to the assembly of a mix-polarity cortex remodelled by actin turnover (naturalistic route for synthetic division),^10,45,54^ the use of this motor protein could render promising results. Alternatively, as Kučera *et al.* showed,^57^ an interesting element that could substitute or supplement myosin activity is anillin, a passive actin crosslinker capable of generating contractile forces. Regardless of the approach, future studies with optimized contractile toolsets could include the MinDE system to efficiently position these actin-based assemblies at mid-cell and deform lipid vesicles.

Collectively, the findings here reported provide a potential new avenue for the equatorial localization of contractile actomyosin rings and the spatial targeting of mechanical forces at the vesicle membrane. Hence, being one step closer to a well-defined self-division of artificial minimal cells, future investigations should consider expanding the system reconstituted here with other division toolsets like a membrane expansion system to engineer membrane growth, an imperative to achieve sustained synthetic division.

## 4. Conclusion

In conclusion, we show that positioning and confinement of contractile actomyosin rings in vesicles to a defined zone of future constriction, ideally in the equatorial region, is possible by employing the versatile MinDE protein system. We thus provide an example of a successful synthetic integration of functional toolkits from different organisms for the division of minimal vesicular systems. As synthetic biology strives to achieve the construction of an artificial cell, minimal modules reconstituted separately must be combined. Working towards the optimization of experimental conditions to meet the needs of all components may be the next challenge of the field. However, the efforts to interlace diverse functional modules might provide the field with new interesting outcomes, like emergent protein functions or unexpected advantageous effects arising from the interplay of completely unrelated families of biomolecules.

## 5. Methods

### Proteins

Actin (alpha-actin skeletal muscle, rabbit), ATTO647-Actin (alpha-actin skeletal muscle, rabbit) and Biotin-Actin (alpha-actin skeletal muscle, rabbit) were purchased from HYPERMOL (Germany). Myosin (rabbit muscle) was purchased from Cytoskeleton Inc (Tebubio GmbH, Offenbach, Germany). Fascin (human, recombinant) was purchased from Cytoskeleton Inc (Tebubio GmbH, Offenbach, Germany) and HYPERMOL (Germany). Stock solutions for all purchased cytoskeletal proteins were obtained by following the handling instructions of the manufacturer. Stock solutions of neutravidin (Thermo Fisher Scientific Inc., Massachusetts, USA, Cat# 31000) were prepared by dissolving the protein in water according to reconstitution instructions.

MinE-His, His-MinD and His-EGFP-MinD were purified as described in previous reports.^58,59^ Briefly, His-tag carrying proteins were purified via affinity chromatography using Ni-NTA columns. Transformed *E.coli* BL21 cells were lysed by sonication and cell lysates were centrifuged to discard debris. Supernatants were loaded into Ni-NTA columns and proteins were eluted in storage buffer (50 mM HEPES, pH 7.25, 150 mM KCl, 0.1 mM EDTA, 1 mM TCEP, 10% Glycerin). Protein purity was confirmed via SDS-PAGE. Protein concentration and protein activity were determined via Bradford assay and ATPase assay, respectively.

### Crowder and density gradient solutions

BSA (Sigma-Aldrich, St. Louis, USA, Cat# A6003) stock solutions were prepared by dissolving the lyophilized powder in Millipore water at approximately 100 g/L as previously described.^9^ To remove undesired debris from BSA solutions, three washing steps were performed with Millipore water using Amicon Ultra-0.5 centrifugal filters 50 kDa MWCO (Merck KGaA, Darmstadt, Germany). Concentration of final stock solutions (ranging from 170 to 300 g/l) were determined via Bradford Assays and stored at -20 °C. Ficoll70 (Sigma-Aldrich, St. Louis, USA, Cat# F2878) was dissolved in Millipore water and left at 4 °C in a rotary shaker for 24 hours. Stock concentrations were calculated from the weight of Ficoll70 added and the final volume of solution obtained (625 g/L). Ficoll70 stocks were subsequently stored at 20 °C. 60% Iodixanol (OptiPrep^TM^, Cat# D1556) was purchased from Sigma-Aldrich (St. Louis, USA).

### Vesicle production

All lipids were purchased from Avanti Polar Lipids (USA). For single-phase vesicles, a lipid-oil emulsion was prepared by dissolving 1-palmitoyl-2-oleoyl-sn-glycero-3phosphocholine (POPC), 1-palmitoyl-2-oleoyl-sn-glycero-3-phospho(1’-rac-glycerol) (POPG) and 1,2-dioleoyl-sn-glycero-3-phosphoethanolamine-N-(cap biotinyl) (sodium salt) (18:1 Biotinyl CAP PE) at a 6.9: 3: 0.1 molar ratio in 2.5 g/L final concentration and drying the mixture for 15 min under a nitrogen stream. To ensure full evaporation of chloroform, mixtures were placed in a vacuum-sealed desiccator for at least 2 hours. Inside a glove box (right before encapsulation experiments) 37.5 µL of decane (TCI Deutschland GmbH, Germany) was added to the dried lipid film and, once dissolved, 1 mL mineral oil (Carl Roth GmbH, Germany) was added and the lipid-oil suspension vigorously vortexed until a clear solution was obtained. For phase-separated vesicles, the lipid mix used was DOPC: DOPG: DPPC: DPPG: Chol (17.5: 7.5: 31.5: 13.5: 30) labelled with 0.001 mol% Atto-655 DOPE binding to the Ld phase. The lipid mix (3.2 mM) was dissolved in chloroform and dried in a glass vial under N2 flow for ∼15 minutes. The dried film was then suspended in a mixture of decane (20 µL) and mineral oil (480 µL) and sonicated at elevated temperatures for ∼30 minutes. Both single single-phase and phase-separated GUVs were produced with the double emulsion transfer method following a recently reported protocol for vesicle generation with purified proteins in 96 well-plates.^60,9^ To ensure iso-osmolar conditions between the inside and outside of GUVs, the osmolarity of inner encapsulating solutions was measured with a osmometer (Fiske Micro-Osmometer model120, Fiske Associates, Norwood, MA, USA) and outer glucose solutions with matching osmolarities were used as outer aqueous environment where GUVs are collected after production for subsequent imaging. The density of the inner mixture depends on the experimental conditions tested but to generate all the vesicles here reported we centrifuged well-plates at 600 × g for 10 min. In the case of single-phase vesicles centrifugation was performed at RT. For phase-separated GUVs, well-plates were centrifuged at 37 °C and the sample was allowed to cool to RT for ∼30 min before imaging.

### Encapsulation of the system in single and double-phase GUVs

Depending on the experimental conditions tested, the final concentrations of the proteins varied but the procedure remained the same. All the steps were performed on ice except for the final encapsulation on 96 well-plates. First, we prepared a 35 µM actin mix (A-Mix) comprised of 86% G-actin, 10% ATTO647-actin and 4% biotinylated actin in water. Once we were ready to encapsulate, we prepared the following inner reaction mix: 4% OptiPrep^TM^, 0.01 g/L Neutravidin, 10 g/L BSA, 10-50 g/L Ficoll70, 3-3.2 MinD (70% His-MinD, 30% EGDP-MinD), 1.6-3 MinE, 1.5-4 µM Actin (from the A-mix), 0.3-2 µM fascin, 0-0.05 µM Myosin and 5mM ATP (frozen stocks supplemented with 5 mM MgCl_2_) in a final buffer concentration of 50 mM KCl, 10 mM Tris-HCl and 5 mM MgCl_2_.

### Fluorescence Microscopy

Imaging of vesicles was performed on a LSM800 confocal laser scanning microscope using a C-Apochromat 40×/1.2 water-immersion objective (Carl Zeiss, Germany). Fluorophores were excited using diode-pumped solid-state lasers: 488 nm (EGFP-MinD) and 640 nm (ATTO647-Actin).

### Image analysis

Processing and analysis of all acquired images were conducted using Fiji (v1.53f51),^61^ custom-written scripts in MATLAB (R2022a) and OriginPro (2021b), the latter also being used for plotting datasets. Z-Stacks were reconstructed in Fiji using the Standard Deviation Z-Projection. For kymographs, the Multi Kymograph function was used and ROIs were drawn manually following the GUVs’ fluorescence intensity at their equatorial cross section with the free-hand selection, fitting them to splines before kymograph retrieval. For aspect ratio calculations, **Fig. 2a,b** GUVs were traced by hand using the free-hand selection and major and minor axes were retrieved from Fiji measurements. For **Fig. 2c**, all time frames from the EGFP channel were processed first with a Gaussian filter of radius 1 and then Otsu-thresholded (frames where GUV contours where incorrectly thresholded were discarded). After filling holes, ROIs retrieved from the particle analyser were fitted to an ellipse to measure the major and minor axes. Intensity profiles in **Fig. 3b** were obtained applying the Plot Profile Fiji function to segmented lines after being fitted to splines. For the waterfall 3D plot in **Supplementary** Fig. 3b, a 5 µm intensity line-plot at the bleb was drawn in Fiji to obtain fluorescence intensity values at six different time points. The data was plotted in Origin using the 3D waterfall plot. In **Fig. 3d**, curvatures were calculated by drawing ROI circles fitting the blebs and retrieving radii measurements from Fiji.

### Analysis and quantification of actomyosin phenotypes inside GUVs

Data were obtained by acquiring Z-stacks of 6 separate experiments (triplicates for presence and absence of Min proteins) after Min oscillation decay for those where Min proteins were added. Vesicles of different sizes with encapsulated actin bundles were classified into four categories (rings, asters, stiff-straight bundles and soft webs) by analysing the acquired Z-stacks and their Standard Deviation Z-Projection. GUV diameters were obtained by manually drawing circle ROIs at the vesicle equator and vesicles were grouped into 4 categories according to their size. GUVs with no actin assemblies inside were not considered for the analysis. Frequencies were calculated from the number of vesicles belonging to each actin-assembly category and the total number of GUVs analysed per condition. Datasets were normalized using the total number of vesicles analysed in each size group.

### Analysis of MinDE-induced folding of actin bundles

To quantify the angle between the two bundle ends attached to the vesicle membrane, the equator of the ATTO647-Actin channel was taken, and this time-series segmented using the “Moments” threshold method. To automatically detect both bundle ends as features and track their position over time, the Fiji plugin ‘TrackMate’ was employed.^62^ The LoG detector was configured with threshold 7, radius 6 px, median filtering, and subpixel localization. Coordinates of the spots detected (two per time frame corresponding to the bundle ends) were obtained with TrackMate’s Simple LAP tracker using the following settings: linking max distance 1 px, gap-closing max distance 15 px, gap-closing max frame gap 1. The angles between the two bundle ends over time were obtained with a MATLAB custom script taking spot coordinates as vector with the vesicle’s centre point as origin.

## Data availability

The data that support the findings of this study are available from the corresponding author upon reasonable request.

## Acknowledgements

The authors would like to thank the MPIB Core Facility for assistance in protein purification, Michaela Schaper for plasmid cloning, Kerstin Röhrl for protein purification, Sandra Ortmeier for lipid preparations and Sigrid Bauer for her advice on vesicle formation and unconditional scientific support. The authors would also like to thank Jan-Hagen Krohn for assistance in confocal microscopy, as well as Adrián Merino-Salomón and Shunshi Kohyama for helpful discussions on crowder conditions and protein encapsulation. This work was supported by the Deutsche Forschungsgemeinschaft (P.S. and M.J.). Y.Q. received funding from the European Union’s Horizon 2020 research and innovation programme under the Marie Skłodowska-Curie grant agreement no. 859416. M.R.-L. and Y.Q. are part of IMPRS-ML and M.R.-L. is part of the ONE MUNICH Project supported by the Federal Ministry of Education and Research (BMBF) as well as the Free State of Bavaria under the Excellence Strategy of the Federal Government and the Länder. The authors would also like to acknowledge the support of the Center for Nanoscience (CeNS), Munich.

## Author Contributions

M.R.-L. and P.S. conceived the study. M.R.-L. designed and performed encapsulation experiments, analyzed data and interpreted results. N.K and M.R.-L. designed and carried out phase-separation experiments. M.R.-L and Y.Q analyzed blebbing vesicles. M.J. provided technical advice on protein conditions for encapsulation. M.R.-L. and P.S. wrote the manuscript and all authors revised and approved the final version of the manuscript.

